# Girls gone wild: Social isolation induces hyperactivity and exploration in aged female mice

**DOI:** 10.1101/2020.04.15.043877

**Authors:** D. Gregory Sullens, Kayla Gilley, Kendall Jensen, Melanie J. Sekeres

**Affiliations:** Department of Psychology and Neuroscience, Baylor University, Waco, Texas, 76798, USA; Department of Biology, Baylor University, Waco, Texas, 76798, USA

**Author notes:** Corresponding author, Address: Department of Psychology Neuroscience, Baylor University, One Bear Place, Waco, Texas, USA, 76798.

**Keywords:** social isolation, aging, female, anxiety, hyperactivity

## Abstract

The likelihood of experiencing social isolation increases later in life, particularly for females. It remains unknown how late-life social isolation impacts cognition and affective behavior in aged mice. We assessed the impact of late-life social isolation in 18-month old female mice. One month of single-housing did not lead to robust depressive-like symptomology, altered social interaction behavior, or sensitivity to context fear acquisition or memory. Rather, isolation increased hyperactivity and exploration, and reduced anxiety-like behavior in the open field and elevated plus maze, findings that have been similarly observed in young female and male mice following early-life isolation. These findings suggest that hyperactivity is a robust behavior following social isolation across the lifespan.

## INTRODUCTION

Given the recent social isolation recommendations in place globally in response to the COVID-19 pandemic, understanding the impact of social isolation on behavior across the lifespan is important for informing interventions that may mitigate the development of mental health disturbances following an isolation period. This is especially critical in aged populations, and particularly in older females, given the increased incidence of living alone in older women.

Rodents, like humans, are social creatures that thrive in grouped-housing conditions. Rodent models have focused on the effects of single-housing isolation-induced stress during critical developmental periods in adolescents and early adulthood (Arakawa, 2018), and typically report increases in depressive-like and anxiety-like behaviors (Berry et al., 2012; Ieraci et al., 2016). Early-life single-housing studies find differential effects of isolation across sex, with females showing less anxiety-like behaviors, and inconsistent memory deficits (Rodgers & Cole, 1993; Bartolomucci et al., 2003; An et al., 2017). Early-life isolation also leads to hyperactivity and increased exploratory behavior, suggesting that isolation may have an anxiolytic effect in young animals (Ashby et al., 2010; Fei et al., 2019). The behavioral impact of social isolation may depend upon the developmental time point of isolation, sex, and duration of isolation.

It remains unknown how late-life social isolation impacts cognition and affective behavior in aged mice. While the likelihood of experiencing social isolation dramatically increases in most aged human populations, this question is of particular relevance to females due to longer life expectancy (Population Reference Bureau). To address the gap in the literature on the effects of social isolation in older animals, we investigated behavioral disturbances in response to late-life isolation in aged female mice.

## METHODS

Female F1 hybrid C57BL/6J (Jackson Labs) × 129S6/SvEvTac (Taconic) served as subjects. Mice were bred in the mouse vivarium at Baylor University. Post-weaning, mice were group-housed (3-5 per cage) in standard shoebox cages with bedding and nesting materials, located in ventilated racks in the rodent vivarium. Throughout the study, mice had *ad libitum* access to food and water, and were maintained on a standard 12hr light-dark cycle (lights on 600hr – 1800hr), with behavioral testing conducted during the light phase of the cycle.

At 18 months of age, home cages of mice were randomly divided into group-housed (n=20 mice) and single-housed social isolation conditions (n=20 mice). For subjects assigned to the social isolation condition, each mouse was individually housed in a standard shoebox cage with bedding and nesting materials, *ad libitum* access to food and water, in ventilated racks in the rodent vivarium for one month. At the end of the one-month isolation period, mice remained in their assigned housing conditions and began a post-rearing behavioral test battery to assess anxiety-like and depressive-like behavior and contextual fear memory (see Figure 1a for experimental timeline).

**Figure 1.**
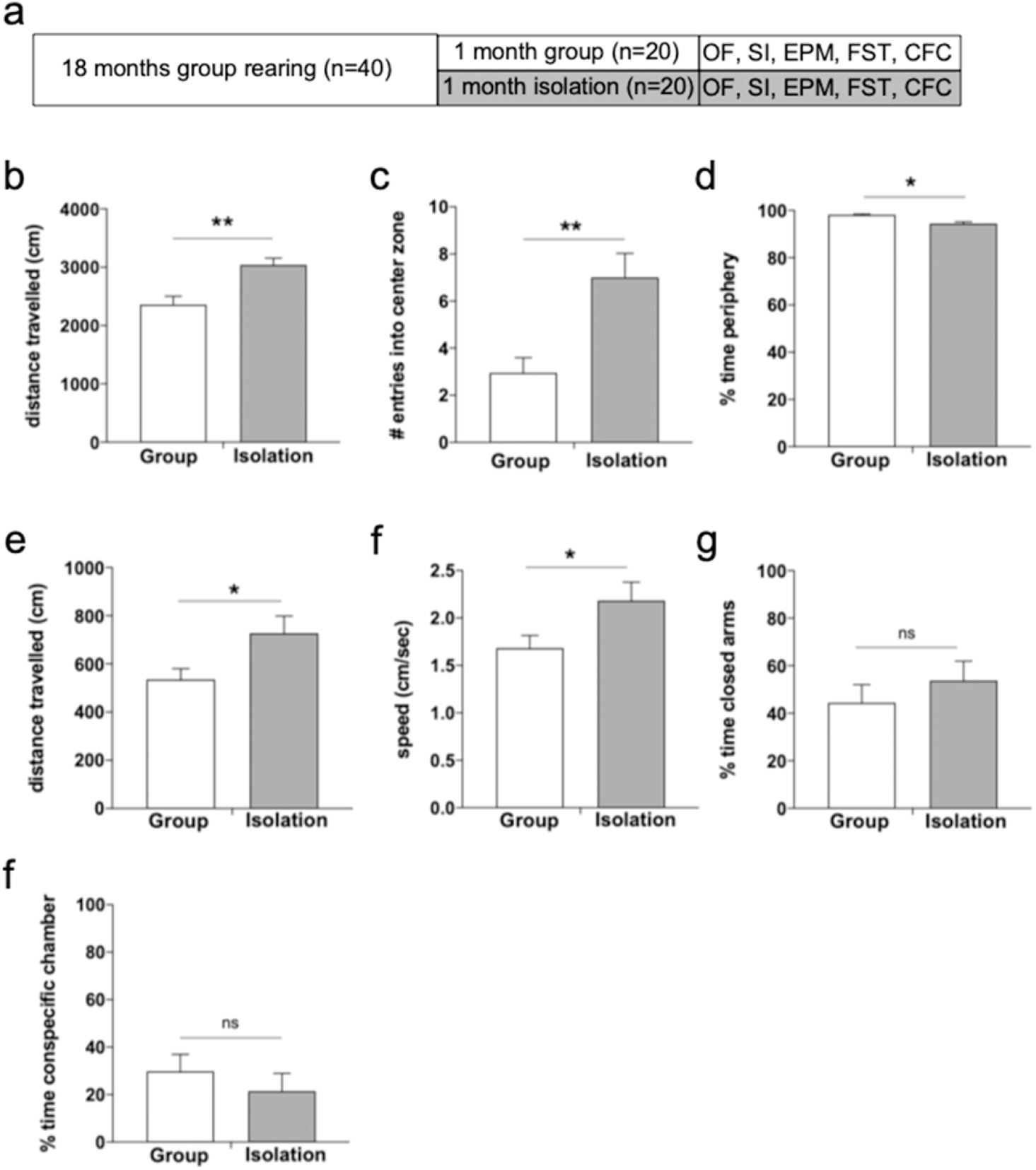
Late-life social isolation increases exploration and hyperactivity in female mice. (a) Experimental timeline. *OF* = open field, *SI* = 3-Chambered Social Interaction Task, *EPM* = Elevated Plus Maze, *FST* = Forced Swim Task, *CFC* = Context Fear Conditioning. In the OF task, (b) isolated mice (grey bars) travel further distance, (c) make more entries into the center of the OF arena, and (d) spend less time in the periphery of the arena than group-housed controls (white bars), indicating a less anxiety-like phenotype. In the EPM task, (d) isolated mice travel further distance and (e) move at a faster speed, (f) but spend an equivalent amount of time in the closed arms of the EPM as controls. (g) Isolated mice spend an equivalent amount of time in the conspecific chamber of the SI arena as controls. Error bars represent the standard error of the mean (SEM). * p < 0.05, ** p < 0.01.

Mice were handled individually by an experimenter for 5 min/day for 5 days prior to beginning behavioral testing. Behavioral tasks were conducted in a least-to-most aversive order, and each test apparatus was cleaned with 70% ethanol between trials. Only a single behavioral task was conducted on each day. *Open Field (OF):* Mice were placed individually in the open field arena (45 cm × 45 cm × 40 cm, white, solid acrylic walls) for 15 min. The percentage of time spent in the different areas (periphery, middle zone, and center zone) of the open field, and the distance and speed travelled were analyzed. Periphery is defined as the area within 10 cm from the edge of the wall. The middle zone is defined as the area 5 cm between the periphery and center zone. The center zone is the 15 cm^2^ area within the middle of the arena. Preference for the periphery of the arena is indicative of an anxiety-like phenotype. *3-Chambered Social Interaction Task (SI)*: The social interaction arena consists of three equally sized clear acrylic chambers (43 cm × 20 cm × 23 cm each chamber) connected by a sliding door on either side of the center chamber. Isolation cages (8 cm diameter, 18 cm height, metal bars spaced 6 mm apart) were located in each of the side chambers. One isolation cage contained a novel sex-matched conspecific mouse. The other cage remained empty during the trial. The test mouse was placed in the center chamber and allowed 5 min to habituate to the chamber. The sliding doors were then raised, the mouse was allowed to freely explore all three chambers of the arena for an additional 5 min. The total time spent exploring the chamber containing the conspecific relative to the other two chambers was measured. Increased time spent in the social side chamber relative to the non-social side chamber indicates a preference for social interaction. *Elevated Plus Maze (EPM):* The white Plexiglas apparatus consisted of two open arms (30 cm long × 5 cm wide × 0 cm high) and two closed arms (30 cm long × 5 cm wide × 15 cm high) extended from a central platform (5 cm × 5 cm) elevated 50 cm above the floor. Mice were individually placed on the central platform facing an open arm and allowed to freely explore the maze for 5 min. The number of entries (all 4 paws) into open and closed arms of the maze, and the total time spent in open and closed parts of the maze were measured. *Forced Swim Task (FST):* A clear Plexiglas cylinder (5 cm diameter, 25 cm high) was filled halfway with room temperature water. The mouse was placed in the chamber for 6 min. The latency to immobility, and the durations of immobility and high activity during the task was calculated. High activity was defined by the SMART behavioral analysis software as ≥ 15% change in pixels between serial video frames during the task. Lower periods of activity are indicative of a learned helplessness phenotype or depressive-like behavior. *Context Fear Conditioning and Testing (CFC):* Fear conditioning was conducted in a chamber (19 cm × 20 cm × 128 cm) with shock-grid floors (bars 3.2 mm in diameter spaced 7.9 mm apart), clear acrylic front and back walls, and aluminum side-walls and roof (Coulborn Instruments). Mice were allowed 2 min to explore the chamber, then received 3 foot shocks (0.5 mA, 2 sec duration, 1 min apart). Mice were removed from the chambers 1 min after the last shock. Twenty-four hours later, the mouse was replaced in the conditioning chamber and freezing behavior was recorded for 3 min. Freezing is a species-specific defense reaction that is typically used as a measure of fear in rodents. Behavior in the chamber was recorded by an overhead camera, and activity levels were analyzed using Freezeframe software (Actimetrix, RRID:SCR_014429).

Activity during other behavioral tasks was recorded by a digital camera and activity levels were analyzed using the SMART video-tracking system (Panlab, RRID: SCR_002852) software. Independent samples t-tests (2-tailed) were conducted for all behavioral tasks. All behavioral statistical analyses were conducted using SPSS 26 (RRID: SCR_002865) by an experimenter blind to experimental condition. A value of *p*□< .05 was considered significant, with figures depicting the mean ± standard error of the mean (SEM). All procedures were approved by Baylor University’s Institutional Care and Use Committee and conducted in accordance with the Guide for the Care and Use of Laboratory Animals from the National Institutes of Health.

## RESULTS

Socially isolated mice engaged in more exploratory behavior in the OF task, travelling a further distance during the OF trial (*t*_(38)_ = −3.70, *p* = 0.001, *d* = 1.169; Figure 1b) than group-housed control mice. Isolated mice displayed a less anxious phenotype than group-housed mice, making more entries into the center of the arena (*t*_(38)_ = −3.33, *p* = 0.002, *d* = 1.054; Figure 1c) and spending less time in the periphery of the arena (*t*_(38)_ = 2.15, *p* = 0.038, *d* = 0.681; Figure 1d).

During the EPM task, Isolated mice explored significantly further distances than group-housed controls (*t*_(38)_ = − 2.29, *p* = 0.028, *d* = 0.723; Figure 1e), and at a greater speed than controls (*t*_(38)_ = −2.12, *p* = 0.041, *d* = 0.700; Figure 1f). The mean percentage of time spent in the in closed arms was not significantly different between group-housed and isolated mice (*t*_(38)_ = −0.85, *p* = 0.403, *d* = 0.268; Figure 1g), suggesting that, despite being more exploratory and hyperactive, late-life isolation did not impact the anxiety-like behavior in the EPM.

During the SI task, surprisingly, no significant difference between groups was observed for the percentage of time spent in the chamber containing the conspecific interaction mouse (*t*_(38)_ = 0.80, *p* = 0.427, *d* = 0.254; Figure 1h), suggesting that the socially isolated mice did not exhibit a preference for social contact, or an aversion to contact, when given the opportunity to interact with a novel conspecific.

In the FST, only a trend was observed in the total time spent immobile during the task (*t*_(38)_ = −1.65, *p* = 0.106, *d* = 0.523; Figure 2a) and the latency to immobility (*t*_(38)_ = −1.49, *p* = 0.146, *d* = 0.582; Figure 2b). Socially isolated mice spent significantly less time performing high activity struggling during the task (*t*_(38)_ = 2.37, *p* = 0.023, *d* = 0.749; Figure 2c), suggesting a mild learned helplessness phenotype during the inescapable swim task.

**Figure 2.**
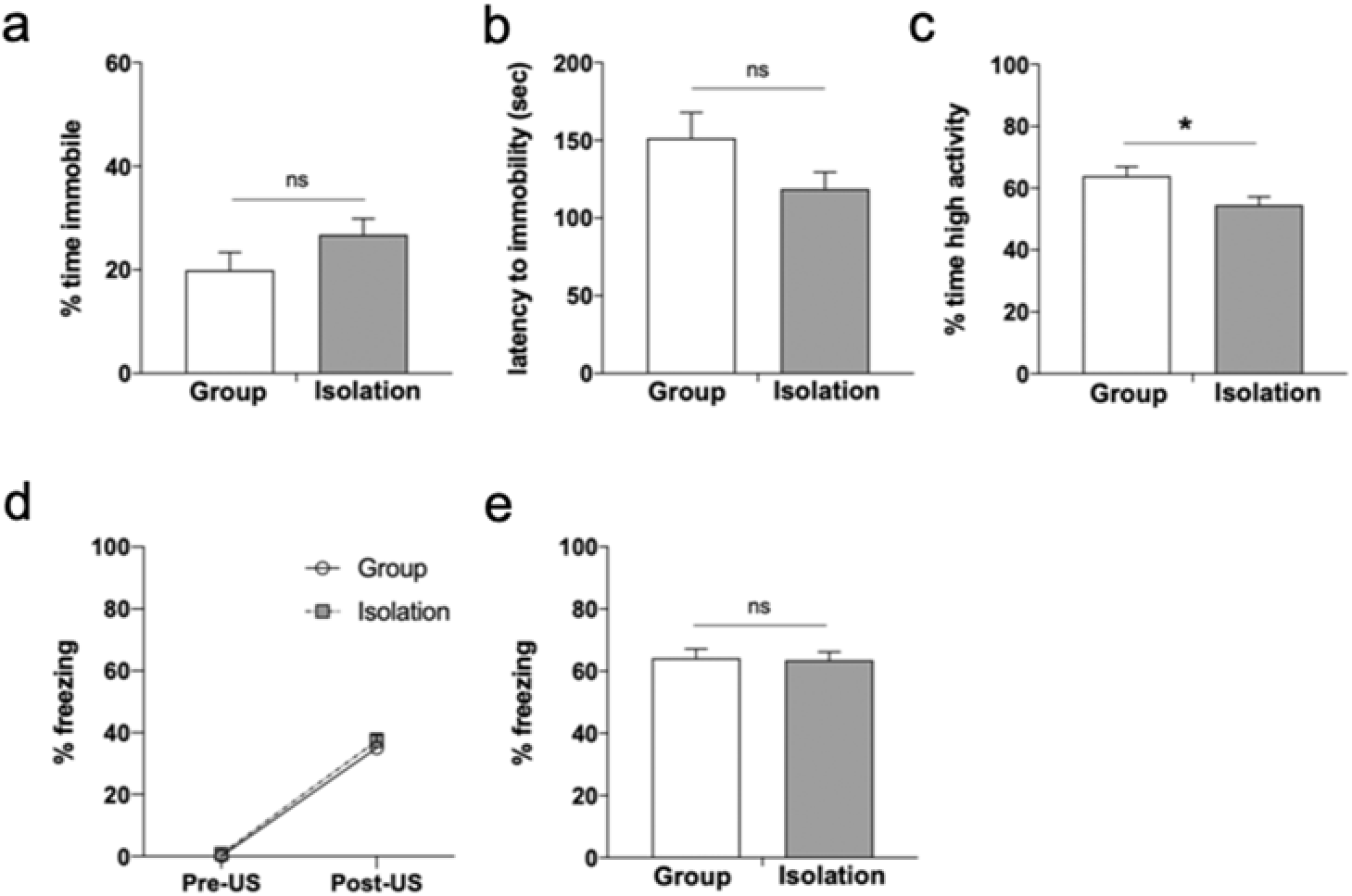
Late-life social isolation does not enhance depressive-like behavior or context fear behavior. (a) Socially isolated mice (grey bars) spend an equivalent percentage of time immobile, and (b) equivalent latency to the first bout of immobility in the FST as group-housed controls (white bars). (c) Isolated mice spend less time performing high-activity struggling behavior in the FST, indicating a mild learned-helplessness phenotype. (d) Isolated and group-housed controls exhibit comparable freezing behavior following the foot-shocks (US) during CFC. (e) Freezing during the context fear test session was equivalent between isolated and group-housed mice, indicating no enhanced fear reactivity or context fear memory following late-life social isolation. Error bars represent the standard error of the mean (SEM). * p<0.05.

Finally, while context fear conditioning is more typically used as a measure of context-dependent memory, it can also be used to assess fear and anxiety-like behavior in rodents. Prior to the presentation of shock (unconditioned stimulus, US), both group-housed and socially isolated mice exhibited comparable levels of activity in the conditioning chamber (*t*_(38)_ = −0.93, *p* = 0.367, *d* = 0.295; Figure 2d), suggesting that late-life isolation alone did not contribute to a baseline increase in anxiety-like behavior during the task. Following the presentation of the three foot-shocks, both groups exhibited equivalent levels of post-US freezing during the conditioning trial (*t*_(38)_ = −0.44, *p* = 0.665, *d* = 0.138; Figure 2d), indicating that isolation did not enhance reactivity to the US. Twenty-four hours later, both groups exhibited comparable levels of freezing during the context memory test (*t*_(38)_ = 0.18, *p* = 0.859, *d* = 0.057; Figure 2e). Together these findings indicate that late-life social isolation did not impair the mouse’s ability to form a context fear memory, nor did it enhance the sensitivity or reactivity of the mice to an aversive shock stimulus.

## DISCUSSION

We assessed exploration and hyperactivity, anxiety-like and depressive-like behavior, and novel context fear memory following social isolation in aged female mice. Isolated mice exhibited increased hyperactivity and exploratory behaviors in the EPM and OF tests relative to controls. Social isolation studies in adolescent or young-adult rodents similarly find isolation increases exploration and locomotor activity, and reduces anxiety-like behavior in the OF, EPM, and nose-poke hole boards (Hilakivi et al., 1989; Ashby et al., 2010; Fei et al., 2019). Observed hyperactivity and exploratory behaviors in novel contexts may be partially a result of declines in GABA-A receptors in the hippocampus and cortex following isolation (Pinna et al., 2006). Young male mice exhibit stronger exploratory behavior in the EPM than females following isolation (Guo et al.,2004), however further studies are needed to determine if these sex differences would be preserved following late-life isolation in a larger sample size. Isolated aged mice displayed less anxiety-like behavior in the OF, travelling into the center of the arena more than group-housed animals. In the EPM, this same risk-taking behavior was not observed, with all mice spending equivalent time in the closed arms of the maze. Previous studies have reported decreased (Guo et al., 2004; Flanigan & Cook, 2011) and comparable (Rodgers & Cole, 1993) anxiety-like behaviors in isolated young mice on the EPM.

Hyperactivity was not observed in an inescapable swim task. Mice showed moderate behavioral despair, as indicated by less time performing high-activity swimming during the FST. No difference in immobility behavior was observed between groups, suggesting that isolation did not induce a robust depressive-like phenotype. The effects of isolation on depressive-like symptoms in young mice are mixed, with reports of more immobility particularly in males (Berry et al., 2012; Takatsu-Coleman, et al., 2013; Liu et al., 2019), while others find less immobility (Kulesskaya et al., 2011) or no difference between single and group-housed animals (Hilvakivi et al., 1989).

Isolation did not impact fear behavior, nor preference for social engagement in aged female mice. The lack of potentiation of a hippocampal-dependent context fear memory is consistent with cellular investigations finding no difference in the electrophysiological properties of CA1 and CA3 following month-long isolation in young mice (Ashby et al., 2010), although isolation did reduce dendritic arborization in CA1 of young mice (Liu et al., 2019), suggesting that isolation stress may structurally remodel the hippocampus. Male mice isolated in young adulthood displayed increased social interaction (Berry et al., 2012) while male rats isolated at weaning displayed reduced interactions and increased freezing behavior (Lukkes et al., 2009).

In summary, late-life social isolation in aged female mice did not lead to robust depressive-like symptomology, altered social interaction behavior, or sensitivity to context fear acquisition and memory. Rather, isolation increased hyperactivity and exploration, findings that have been similarly observed in both sexes following early-life isolation. In human populations, social isolation and loneliness correlate with cognitive decline, depression, and anxiety (Cacioppo et al., 2006; Domènech-Abella et al., 2019). Understanding the behavioral impacts of late-life isolation is especially relevant given the current self-isolation recommendations, particularly for females whose are more likely than males to live alone. By identifying behavioral disturbances in behavior following isolation the results will be useful for informing the likely cognitive and psychological impact of isolation in response to the recent social isolation experiences of the COVID-19 pandemic.

## ACKNOWLEDGEMENTS

We thank Lee Lowe and Natashia Howard for technical contributions.

## CONTRIBUTIONS

D.G. Sullens and M.J. Sekeres developed the study concept and design. Testing and data collection were performed by D.G. Sullens, K. Gilley, and K. Jensen. Data analysis and interpretation was performed by M.J. Sekeres. D.G. Sullens and M.J. Sekeres drafted the manuscript. All authors contributed to manuscript review and revisions. All authors approved of the final version of the manuscript.

## OPEN PRACTCES DATA SHARING STATEMENT

The experiment reported in this article was not formally pre-registered. Neither the data nor the materials have been made available on a permanent third-party archive. Requests for the data or materials can be sent via email to the corresponding author at.

## REFERENCES

An, D., Chen, W., Yu, D. Q., Wang, S. W., Yu, W. Z., Xu, H., Wang, D. M., Zhao, D., Sun, Y. P., Wu, J. C., Tang, Y. Y., & Yin, S. M. (2017). Effects of social isolation, re-socialization and age on cognitive and aggressive behaviors of Kunming mice and BALB/c mice. Animal Science Journal, 88(5), 798–806. https://doi.org/10.1111/asj.12688

Arakawa, H. (2018). Ethological approach to social isolation effects in behavioral studies of laboratory rodents. Behavioural Brain Research, 341, 98–108. https://doi.org/10.1016/j.bbr.2017.12.022

Ashby, D. M., Habib, D., Dringenberg, H. C., Reynolds, J. N., & Beninger, R. J. (2010). Subchronic MK-801 treatment and post-weaning social isolation in rats: Differential effects on locomotor activity and hippocampal long-term potentiation. Behavioural brain research, 212(1), 64–70.

Bartolomucci, A., Palanza, P., Sacerdote, P., Ceresini, G., Chirieleison, A., Panerai, A. E., & Parmigiani, S. (2003). Individual housing induces altered immunoendocrine responses to psychological stress in male mice. Psychoneuroendocrinology, 28, 540–558. https://doi.org/10.1016/S0306-4530(02)00039-2

Berry, A., Bellisario, V., Capoccia, S., Tirassa, P., Calza, A., Alleva, E., & Cirulli, F. (2012). Social deprivation stress is a triggering factor for the emergence of anxiety- and depression-like behaviours and leads to reduced brain BDNF levels in C57BL/6J mice. Psychoneuroendocrinology, 37(6), 762–772. https://doi.org10.1016/j.psyneuen.2011.09.007

Cacioppo, J. T., Hughes, M. E., Waite, L. J., Hawkley, L. C., & Thisted, R. A. (2006). Loneliness as a specific risk factor for depressive symptoms: Cross-sectional and longitudinal analyses. Psychology and Aging, 21(1), 140–151. https://doi.org/10.1037/0882-7974.21.1.140

Domènech-Abella, J., Mundó, J., Haro, J. M., & Rubio-Valera, M. (2019). Anxiety, depression, loneliness and social network in the elderly: Longitudinal associations from The Irish Longitudinal Study on Ageing (TILDA). Journal of Affective Disorders, 246 (November 2018), 82–88. https://doi.org/10.1016/j.jad.2018.12.043

Fei, X. Y., Liu, S., Sun, Y. H., & Cheng, L. (2019). Social isolation improves the performance of rodents in a novel cognitive flexibility task. Frontiers in Zoology, 16(1), 43.

Flanigan, T. J., & Cook, M. N. (2011). Effects of an early handling-like procedure and individual housing on anxiety-like behavior in adult C57BL/6J and DBA/2J mice. PLos One, 6(4), Article: e19058. doi:10.1371/journal.pone.0019058

Guo, M., Wu, C. F., Liu, W., Yang, J. Y., & Chen, D. (2004). Sex difference in psychological behavior changes induced by long-term social isolation in mice. Progress in Neuro-Psychopharmacology and Biological Psychiatry, 28(1), 115–121. https://doi.org/10.1016/j.pnpbp.2003.09.027

Hilakivi, L. A., Ota, M., & Lister, R. G. (1989). Effect of isolation on brain monoamines and the behavior of mice in tests of exploration, locomotion, anxiety and behavioral ‘despair’. Pharmacology Biochemistry & Behavior, 33(2), 371–374. https://doi.org/10.1016/0091-3057(89)90516-9

Ieraci, A., Mallei, A., & Popoli, M. (2016). Social isolation stress induces anxious-depressive-like behavior and alterations of neuroplasticity-related genes in adult male mice. Neural Plasticity, 2016. https://doi.org/10.1155/2016/6212983

Kulesskaya N., Rauvala H., & Voikar V. (2011). Evaluation of social and physical enrichment in modulation of behavioural phenotype in C57BL/6J female mice. PLoS ONE, 6(9), Article: e24755. doi:10.1371/journal.pone.0024755

Lukkes, J. L., Mokin, M. V., Scholl, J. L., & Foster, G. L. (2009). Adult rats exposed to early-life social isolation exhibit increased anxiety and conditioned fear behavior, and altered hormonal stress responses. Hormones and Behavior, 55(1), 248–256. doi:10.1016/j.yhbeh.2008.10.014

Liu, N., Wang, Y., An, A. Y., Banker, C., Qian, Y. H., & O’Donnell, J. M. (2019). Single housing‐induced effects on cognitive impairment and depression‐like behavior in male and female mice involve neuroplasticity‐related signaling. European Journal of Neuroscience. Advanced online August 31, 2019. doi: 10.1111/ejn.14565

Pinna, G., Agis-Balboa, R. C., Zhubi, A., Matsumoto, K., Grayson, D. R., Costa, E., & Guidotti, A. (2006). Imidazenil and diazepam increase locomotor activity in mice exposed to protracted social isolation. Proceedings of the National Academy of Sciences, 103(11), 4275–4280.

Population Reference Bureau. (n.d.). International data. PRB. https://www.prb.org/international/geography/world

Rodgers, R. J., & Cole, J. C. (1993). Influence of social isolation, gender, strain, and prior novelty on plus-maze behaviour in mice. Physiology & Behavior, 54(4), 729–736. https://doi.org/10.1016/0031-9384(93)90084-S

Takatsu-Coleman, A. L., Patti, C. L., Zani, K. A., Zager, A., Carvalho, R. C., Borcoi, A. R., Ceccon, L. M. B., Berros, L. F., Tufik, S., Anderson, M. L., & Frussa-Filho, R. (2013). Short-term social isolation induces depressive-like behaviour and reinstates the retrieval of an aversive task: Mood-congruent memory in male mice? Journal of Psychiatry & Neuroscience, 38(4), 259–268. doi: 10.1503/jpn.120050z

